# Domestic dogs in indigenous Amazonian communities: key players in *Leptospira* cycling and transmission?

**DOI:** 10.1101/2023.09.19.558554

**Authors:** Diego A. Guzmán, Eduardo Diaz, Carolina Sáenz, Hernán Álvarez, Rubén Cueva, Galo Zapata-Ríos, Belé Prado-Vivar, Mercy Falconí, Talima Pearson, Verónica Barragán

## Abstract

**Background:** Leptospirosis is the world’s most common zoonotic disease. Mitigation and control rely on pathogen identification and understanding the roles of potential reservoirs in cycling and transmission. Underreporting and misdiagnosis obscure the magnitude of the problem and confound efforts to understand key epidemiological components. Difficulties in culturing hamper the use of serological diagnostics and delay the development of DNA detection methods. As a result, especially in complex ecosystems, we know very little about the importance of different mammalian host species in cycling and transmission to humans.

**Methodology/Principal Findings:** We sampled five indigenous Kichwa communities living in the Yasuní National Park in the Ecuadorian Amazon basin. Blood and urine samples from domestic dogs were collected to assess the exposure of these animals to *Leptospira*, and to identify the circulating species. Microscopic Agglutination Tests with a panel of 22 different serovars showed anti-leptospira antibodies in 36 sampled dogs (75%), and 10 serotypes were detected. Two DNA-based detection assays revealed pathogenic *Leptospira* DNA in 18 of 19 dog urine samples (94.7%). Amplicon sequencing and phylogenetic analysis of 16s rDNA and SecY genes from 15 urine samples revealed genetic diversity within two of three different *Leptospira* species: *noguchii* (n=7), *santarosai* (n=7), and *interrogans* (n=1).

**Conclusions/Significance:** The high prevalence of antibodies and *Leptospira* DNA provides strong evidence for high rates of past and current infections. Such high prevalence has not been previously reported for dogs. These dogs live in the peridomestic environment in close contact with humans, yet they are free-ranging animals that interact with wildlife. This complex web of interactions may explain the diverse types of pathogenic *Leptospira* observed in this study. Our results suggest that domestic dogs are likely to play an important role in the cycling and transmission of *Leptospira*. Future studies in areas with complex ecoepidemiology will enable better parsing of the significance of genotypic, environmental, and host characteristics.

**Author Summary:** People around the world interact with a wide range of animals, but one of the closest is the domestic dog. Dogs can be reservoirs of several zoonotic infectious diseases, including leptospirosis. The frequent ecological interactions between people, dogs and wildlife in indigenous communities living in the Amazon basin might increase the complexity of leptospirosis transmission, in comparison with what has been described for other settings. In the Amazon basin, wild animals and domestic animals may act as reservoirs of the pathogen, excreting the bacteria through their urine. In this work we analyzed serum and urine samples from dogs living with Kichwa communities from the Yasuní National Park in Ecuador. Serum samples were analyzed with MAT and urine samples with a qPCR that detect the presence of pathogenic *Leptospira*. Our results suggest that a high percentage of dogs are shedding the pathogen and we identified three different *Leptospira* species. Our serological analysis suggests the presence of ten serovars in dogs and their high exposure to *Leptospira*. These findings provide important insights into the epidemiology of leptospirosis in this ecosystem, suggesting that dogs are likely to play a critical role in the transmission of the disease.

## Introduction

Leptospirosis is a zoonosis that continues to be an important, albeit neglected infectious disease affecting humans and animals worldwide [1]. Leptospirosis particularly affects people living in poverty and with poor sanitation [2–5]. Humans contract leptospirosis when injured skin or mucous membranes are exposed to pathogenic *Leptospira spp.* bacteria by eating contaminated meat, direct contact with urine, or contact with urine-contaminated environments [6–8]. Mild disease occurs in most cases, characterized by a nonspecific, febrile, flu-like illness. On the other hand, severe leptospirosis can cause dysfunction of the kidneys, lungs, and liver, leading to the death of the patient in approximately 5% of cases [9,10]. Unfortunately, diverse symptoms and co-circulation of other febrile diseases (e.g., Dengue and Malaria) leads to under- or misdiagnosis, resulting in an underestimation of the prevalence and severe epidemiological knowledge gaps in our understanding of reservoirs and modes of transmission [7,11,12].

A diverse array of domestic and wild animals likely serve as reservoirs by excreting *Leptospira* in their urine [13–15]. However, like humans, we know little about the prevalence and shedding of these pathogens in peridomestic animal hosts in rural settings [15–20], and reports indicating the seroprevalence of the pathogen in wild animals are even more scarce [21–27]. These important epidemiological parameters may be similar across regions, but site-specific attributes such as host densities and interactions (with human and non-human hosts) are likely highly important for the circulation and transmission of *Leptospira* [16]. For effective control and prevention of leptospirosis, it is vital to explore how local factors influence disease epidemiology.

Humans across the world interact with a variety of animals, but one of the closest is the domestic dog. The exposure of dogs to pathogenic leptospires may depend on: 1) interactions with, or proximity to other domestic and wild animals, and 2) attributes of the local environment in which the animals live, such as hygiene and local weather conditions [28–31]. Dogs can present with mild to severe signs similar to humans; the most common signs are anorexia, lethargy, diarrhea, jaundice, fever, and weakness. Severe leptospirosis in dogs can progress to kidney failure, liver failure, shock, and often death [32]. Importantly, dogs can excrete the bacteria in their urine even without showing signs of infection [33–36]. While the role of dogs in leptospirosis transmission remains poorly understood, particularly in rural communities, their proximity to humans and potential to excrete the pathogen in the peridomestic environment suggests that these animals may play an important role in the epidemiology of human disease [29,37–43].

Kichwa people, one of the indigenous ethnic groups that live in the Ecuadorian Amazon basin, own domestic dogs. However, unlike dogs in most urban settings, these animals are not confined indoors or the immediate vicinity of the homes; they drink water from the river or stagnant pools, roam freely in the forest, and hunt small wild animals [38,44]. This behavior, combined with the region’s climatic conditions which favors the environmental persistence of *Leptospira*, increases the likelihood of exposure and transmission of pathogenic *Leptospira* species to humans and wildlife. Thus, the proximity of dogs to people [44], especially children, may be an important risk factor for leptospirosis in these indigenous communities.

In this study, we investigated whether domestic dogs from five Kichwa communities living on the riverbanks of the Napo River in the Ecuadorian Amazon were exposed to leptospirosis. In addition, we aimed to determine if these dogs were shedding leptospires by detecting DNA of the pathogen in their urine and thus exposing their owners to leptospirosis. Our results suggest that domestic dogs likely play an important role in the local epidemiology of leptospirosis.

## Materials and Methods

### Study Site

This study was carried out in the northern part of Yasuní National Park, in eastern Ecuador (00°30’14’’ S; 76°28’19’’ W). This protected area contains parts of the territory of five Kichwa indigenous communities (approximately 88 000 ha). Yasuní National Park is located on the western Amazon and has been recognized as a major biodiversity hotspot [45]. Encompassing almost 1 million ha, Yasuní is one of the most species diverse forests in the world [46,47]. The study area is classified as a tropical rainforest [48], dominated by large tracts of *terra firme* forest mixed with smaller extensions of palm swamps. There are still large expanses of continuous undisturbed vegetation in the eastern and southern portions of the park, but its northern and western boundaries are surrounded by a growing matrix of pastures, agricultural lands, and secondary vegetation [49].

With a population of more than 100,000 people, the Kichwa constitutes Amazonian Ecuador’s largest indigenous group [50]. This study was performed in five Kichwa communities (Pompeya, Indillama, Nueva Providencia, Sani Isla y San Roque) located along the Napo River and established 40 years ago when a few families moved to the area looking for new hunting grounds (Fig 1). Although some ecotourism activities occur in the area, the local economy is primarily based on subsistence agriculture (mainly plantain and manioc), and a high percentage of protein intake comes from wild meat.

**Fig 1.**
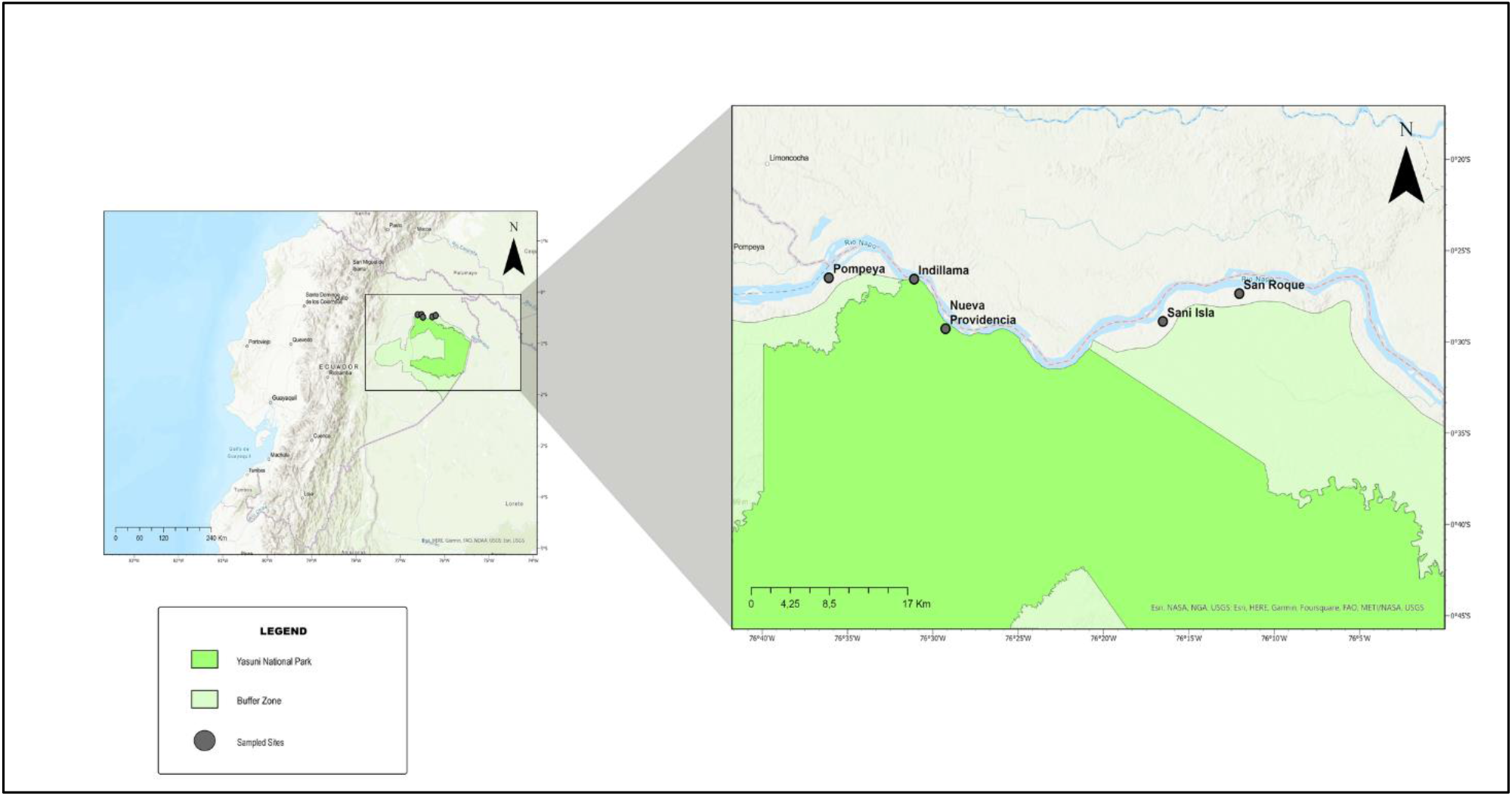
Geographic location of study sites in the Yasuní Biosphere Reserve. The five Kichwa communities that were sampled are shown as grey dots: Pompeya (lat:-0.44148, long:-76.60161), Indillama (lat:-0.44281, long:-76.5184), Nueva Providencia (lat:-0.48804, long:-76.48771), Sani Isla (lat:-0.48152, long:-76.27553), and San Roque (lat:-0.45611, long:-76.20086). The intense green color shows the area that is defined for the Yasuní National Park, and the lighter green color indicates the park buffer zone.

### Sampled population

During June and November 2018, a dog census was performed by the Wildlife Conservation Society – Ecuador (WCS – Ecuador). The aim of this census was to serve as a baseline for population trends and a health surveillance program in the five Kichwa communities in northern Yasuní National Park (Pompeya, Indillama, Nueva Providencia, Sani Isla, and San Roque). It was carried out by a local biologist to assess the number of dogs owned by each household. The percentage of households with dogs in the census varied between communities (Pompeya 50%, Indillama 100%, Nueva Providencia 50%, Sani Isla 76%, and San Roque 91%). The total domestic dog population was 550 dogs, and approximately 10% of the dogs (n=51) were sampled to test for various infectious diseases.

### Sample collection

Ethical approval for sample collection was issued by CEIDA-USFQ (2017-010). Urine and blood samples were collected in August 2019 from domestic dogs living in the communities as follows. Blood samples we collected from the cephalic vein for all domestic dogs except for three, resulting in a total sample size of 48 dogs: Pompeya (n=7), Sani Isla (n=12), San Roque (n=12), Indillama (n=5), and Nueva Providencia (n=12). We collected urine samples from 19 male dogs by transurethral catheterization. It was not possible to get this type of sample from all dogs due to the absence of urine in the bladder of some animals. Transurethral catheterization is difficult in female dogs and thus urine samples from female dogs were not collected. To prevent DNA degradation, 4 ml of urine was added to 4 ml of 2X DNA/RNA Shield_®_ (Zymo, USA). Serum and urine samples were stored and transported on ice to the Institute of Microbiology at Universidad San Francisco de Quito and thereafter maintained at −20°C until DNA extraction.

### Molecular detection of *Leptospira*

The 48 dog serum samples were analyzed by the National Reference Laboratory for Animal Diagnostics (AGROCALIDAD) using Microscopic Agglutination Test (MAT) performed with a panel of 22 different available serovars (Bratislava, Autumnalis, Icterohaemorrhagiae, Canicola, Hardjo, Grippotyphosa, Wolffi, Saxkoebing, Shermani, Celledonis, Javanica, Tarassovi, Pyrogenes, Australis, Bataviae, Andamana, Castellonis, Sejroe, Copenhageni, Pomona, Hebdomadis, Djasiman). Reactive samples with titers ≥ 1:100 were considered positive for detecting anti-leptospiral antibodies. MAT results were visualized by dark field microscopy, and the final titer was assigned as the serum dilution that promotes 50% of agglutination. The serovar with the highest titer was recorded in samples that reacted with multiple serovars. If a sample reacted with more than one serovar with the same titers and without showing a unique highest titer, samples were labeled as “cross-reactive”. Characterized local strains are not available in Ecuador, therefore the serology is performed with non-local strains and cross-reaction results are common.

Molecular detection of pathogenic *Leptospira* spp. was performed on the 19 dog urine samples. Samples were thawed on ice and centrifuged at 4500 × g for 20 min at 4°C. The supernatant was discarded, and 200 μL of the pellet material was used for DNA extraction using DNeasy Blood and Tissue kit (Qiagen, CA, USA). DNA was stored at –20°C. Two previously described Taqman assays were used for molecular detection of pathogenic *Leptospira*: one assay targets the *lipL32* gene [51] and the other, SNP 111, targets a SNP in the *16S rDNA* gene of pathogenic *Leptospira* [17]. A sample was considered positive when at least one of the assays (*lipL32* or SNP 111) detected leptospiral DNA. The redundancy of two independent assays reduces the likelihood of false negatives that, in our experience, is relatively common and due to the large genetic diversity of pathogenic *Leptospira.* Using 2 assays also minimizes false negatives by mitigating against the inherent stochasticity of capturing a PCR target in low quantity samples.

### Amplicon sequencing for *Leptospira* species identification

Primers rrs2 and SecYIV, described by Ahmed et al. [52,53], were used to amplify a fragment of the *16s rDNA* and *SecY* genes from positive samples. Amplicons were sequenced using Oxford Nanopore Technologies. PCR amplicons were obtained using the Q5 High-Fidelity Master Mix (New England, BioLabs), 0.4 μM each primer, and 2.5 μL of DNA template in a final reaction volume of 25 μL. PCR protocol consisted of an initial step at 98°C for 30 seconds, followed by 30 cycles of 10 s at 98°C, 30 s at 58°C or 54°C for rrs2 and SecYIV, respectively, and 30 s at 72°C, followed by final extension step for 2 minutes at 72°C. Amplicons were purified using AMPure XP magnetic beads (Beckman Coulter, USA) following manufacturer instructions and then quantified in a Qubit 3.0 fluorometer (Thermo Fisher Scientific) using the Qubit™ 1X dsDNA, high sensitivity kit (Thermo Scientific, Invitrogen, USA). The quantified samples were normalized to a concentration of 3,0 ng/uL and sequenced following the Oxford Nanopore Library preparation protocol of the Ligation sequencing kit (SQK-LSK109) (Oxford Nanopore Technologies, UK). Finally, 5,72 ng of the library was loaded into a MinION flow cell (FLO-MIN 106). Most reads were obtained during the first 12 hours of the run. Reads were basecalled and demultiplexed using the Guppy software (version 3.4.5) (Oxford Nanopore Technologies, UK) [54] and Porechop (version 0.2.4) (https://github.com/rrwick/Porechop) respectively.

### Sequence analysis for *Leptospira* species identification

*Leptospira* sequences were initially screened using BLASTn command line software (version 2.9.0-2) [55]. This step was implemented to filter out non-*Leptospira* reads obtained during sequencing. Then, sequences were aligned with minimap2 (version 2.22) [56] and visualized in Tablet (version 1.21.02.08) [57]. *L. interrogans* serovar Copenhageni FioCruz L1-130 chromosome 1 (NC_005823.1) was used as a reference genome for the alignment. All the reads that were mapped to the corresponding genes (*16s rDNA* and *SecY*) were filtered to a new file and aligned in MEGA-X (version 10.1.8) [58]. The consensus sequences were obtained using the EMBOSS cons online tool https://www.ebi.ac.uk/Tools/msa/emboss_cons/ [59]. Sequences from both genes were concatenated and compared with representative sequences of each species of *Leptospira* obtained from GenBank. A phylogenetic tree was built in MEGA-X using the Neighbor-Joining method [60], with the Maximum Composite Likelihood model [61] and 500 bootstraps. Finally, the phylogenetic tree was visualized using iTOL [62].

All raw sequence reads were deposited in the NCBI’s Sequence Read Archive (SRA) under Bioproject Number PRJNA758395, and SRA accession numbers SRX12007895-SRX12007909.

## Results

### High seroprevalence among domestic dogs from Kichwa communities

Anti-leptospira antibodies were registered in 36 of the 48 dogs (75% - CI: 60.4-86.4) with titers ≥1:100. Tarassovi serovar was the most predominant (n=5) followed by Australis (n=3), Pyrogenes (n=3), Saxkoebing (n=2), Wolffi (n=2), Canicola (n=2), Hardjo (n=1), Grippotyphosa (n=1), Sejroe (n=1) and Shermani (n=1). 15 samples cross-reacted with more than one serovar. MAT titers for each sample are detailed in Table S1. Dogs from the San Roque community showed the highest positivity (92%, n=11/12), Nueva Providencia (83%, n=10/12), Indillama (80%, n=4/5), Sani Isla (58%, n=7/12) and Pompeya (57%, n=4/7) communities (Table 1).

**Table 1.** *Leptospira* serovars detected in domestic dogs living in Kichwa communities.

| Communities | Seropositivity (%) | Cross-reaction (n) | Predominant Serovars (n) |  |  |  |  |  |  |  |  |  |
| --- | --- | --- | --- | --- | --- | --- | --- | --- | --- | --- | --- | --- |
|  |  |  | Har | Aus | Tar | Wol | Gri | Can | Sax | She | Ser | Pyr |
| San Roque | 92% | 4 | 1 | 2 | 2 | - | - | - | 1 | - | - | 1 |
| Nueva Providencia | 83% | 5 | - | 1 | 1 | - | - | - | - | - | 1 | 2 |
| Indillama | 80% | 2 | - | - | 1 | - | - | 1 | - | - | - | - |
| Sani Isla | 58% | 2 | - | - | - | 2 | 1 | 1 | 1 | - | - | - |
| Pompeya | 57% | 2 | - | - | 1 | - | - | - | - | 1 | - | - |

CR=Cross-reaction; Har=Hardjo; Aus=Australis; Tar=Tarassovi; Wol=Wolffi, Gri=Grippotyphosa; Can=Canicola; Sax=Saxkoebing; She=Shermani; Sej=Sejroe; Pyr=Pyrogenes.

### Detection of pathogenic *Leptospira* DNA in a high percentage of dogs

*Leptospira* DNA was detected in 18 of 19 dog urine samples (97.4% - CI: 73.9-99.8). Some samples that were negative for *lipL32* were positive for SNP111 and vice-versa (Table 2). The SNP111 assay detected *Leptospira* DNA in 7 samples that the *lipL32* assay did not detect. Likewise, the *lipL32* assay detected *Leptospira* DNA in 8 samples that the SNP111 assay did not detect. Only three samples were positive for both assays. Further sequencing analysis confirmed the presence of pathogenic leptospira in most samples. Interestingly, serum samples from only eight out of these 18 PCR-positive dogs registered MAT titers ≥1:100 (Table 2).

**Table 2.** *Leptospira* seropositivity, and detection and identification of pathogenic *Leptospira* DNA in dog urine samples.

| Individuals | Origin | *lipL32*<br>gene | SNP 111 | MAT<sup>1</sup> | Predominant<br>Serovar | *Leptospira* spp.<br>identified |
| --- | --- | --- | --- | --- | --- | --- |
| M02 | Pompeya | N | P | NA | - | *L. noguchii* |
| M03 | Pompeya | N | P | 100 | Tarassovi | *L. interrogans* |
| M08 | Pompeya | N | P | NA | - | *L. noguchii* |
| M10 | Pompeya | P | P | NA | - | *L. noguchii* |
| M14 | San Roque | P | N | 400 | Australis | *L. noguchii* |
| M18 | San Roque | P | N | N | - | NS |
| M19 | San Roque | P | N | 100 | Pyrogenes | *L. santarosai* |
| M21 | San Roque | P | N | 100 | Cross-reaction<sup>2</sup> | *L. santarosai* |
| M23 | Sani Isla | P | N | 400 | Wolffi | *L. santarosai* |
| M27 | Sani Isla | N | P | N | - | *L. noguchii* |
| M28 | Sani Isla | P | N | 200 | Canicola | *L. noguchii* |
| M31 | Sani Isla | P | N | 100 | Cross-reaction | NS |
| M34 | Sani Isla | N | P | 100 | Saxkoebing | *L. santarosai* |
| M38 | Nueva<br>Providencia | P | P | 100 | Cross-reaction | *L. santarosai* |
| M44 | Nueva<br>Providencia | N | P | 200 | Tarassovi | *L. santarosai* |
| M47 | Indillama | N | N | N | - | - |
| M49 | Indillama | N | P | 100 | Cross-reaction | *L. noguchii* |
| M51 | Indillama | P | N | 100 | Canicola | NS |

| M53 | Pompeya | P | P | 200 | Shermani | *L. santarosai* |
| --- | --- | --- | --- | --- | --- | --- |

N= negative; P= positive; NA= blood sample not available; NS= not able to be sequenced.
<sup>1</sup> When multiple serovars were reactive, only the one with the highest titers was reported.
<sup>2</sup> When multiple serovars with the same titers were reactive, it was reported as “Cross-reaction.”

Three out of 18 samples did not yield amplicon qualities sufficient for sequencing. Three species of *Leptospira* were identified from 15 urine samples by sequencing 541 bp and 202 bp fragments of the *16S rDNA* and *SecY* genes, respectively. After concatenation, a fragment of approximately 700 bp was analyzed for species identification. *Leptospira noguchii* (n=7) was present in at least one sample from each community except for Nueva Providencia, *Leptospira santarosai* (n=7) was detected in all communities except for Indillama, and *Leptospira interrogans* was detected in a single sample from Pompeya (Fig 2).

**Fig 2.**
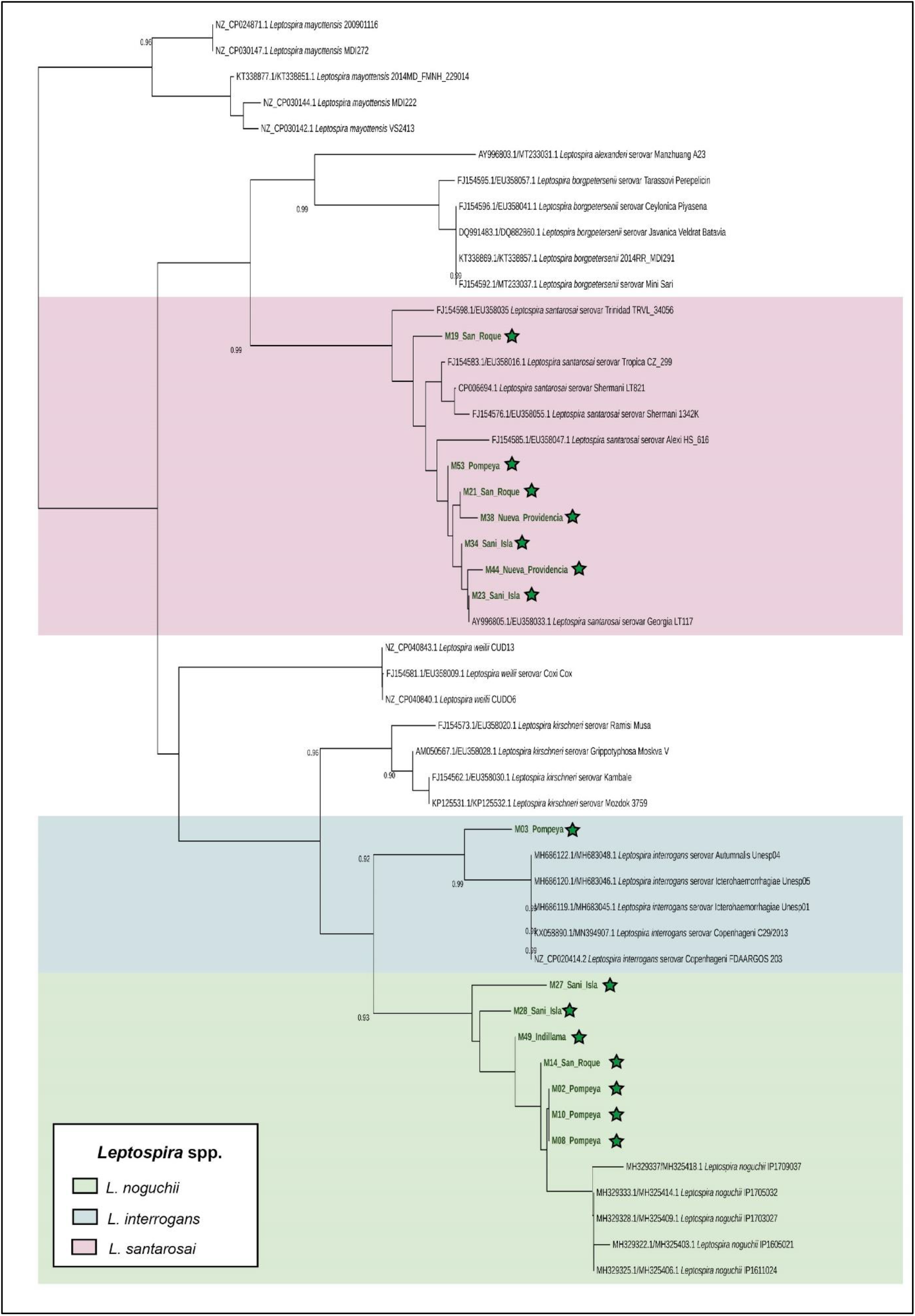
Molecular phylogenetic analysis of a 466 bp fragment obtained by concatenating a 266 bp fragment of the *16S rDNA* and a 200 bp fragment of the *SecY* gene. Bootstrap values (500 replicates) > 0.90 are displayed. The tree was rooted with sequences from *Leptonema illini* DSM 21528 (not depicted on the tree). The samples sequenced from this project are indicated by green stars.

## Discussion

The Amazon basin would appear to be an ideal environment for the circulation of pathogenic *Leptospira* species due to its tropical climate, year-round rainfall, and the high diversity of wildlife that can act as reservoirs. However, very few cases of human leptospirosis are reported in the Ecuadorian Amazon basin, with approximately 38 patients annually from 2015 to 2022 [63]. These few reported cases undoubtedly underestimate prevalence; leptospirosis has been neglected in this region for decades and recently, the public health system has been completely focused on the COVID-19 pandemic [64]. Moreover, public health clinics are challenging to access due to long distances from communities. For communities like Sani Isla and San Roque, the nearest health centers are Añangu (20 km away) and El Eden (18.36 km away), respectively, entailing traveling long distances by foot and boat. For this reason, medical brigades rarely visit these communities. Lack of resources, government priorities, and the remoteness of this area make it difficult to accurately determine the prevalence of diseases, including leptospirosis.

Human interaction with infected animals is a significant risk factor for leptospirosis. Dogs, in particular, live in the peridomestic environment and interact closely with all members of the human community. At the same time, domestic dogs interact with wildlife in the adjacent forest and environment by influencing activity patterns, reducing abundance through direct predation and harassment, and generally disturbing ecosystems [65–67]. Based on the 2018 census, 550 dogs lived in the five participating Kichwa communities. These dogs roam freely in search of food and water, potentially exposing them to leptospirosis and other zoonotic diseases (e.g., rabies, canine distemper, and parvovirus) circulating among wildlife [65,68,69]. Unfortunately, dog vaccinations, deworming, and sterilization in Kichwa communities are not routinely performed due to accessibility difficulties. The prevalence, behavior, and lack of veterinary care of domestic dogs therefore increases the risk of zoonotic disease transmission among domestic animals, wildlife, and humans.

Our serological results in 51 dogs from a population of ∼550 individuals suggests that a high percentage of dogs have been exposed to pathogenic *Leptospira*. The serovars Canicola and Pyrogenes were present in only two and three dogs (respectively), which was not expected because elsewhere, these are the most common serovars in dogs [13,70–73] and are suspected to be responsible for transmission between dogs and humans [13,38,43,72,74]. Additionally, several serovars found in the dogs of our study, such as Tarassovi and Australis, have been primarily associated with pigs, cows, and other small mammals [13,22,70–72,74–77]. The diversity of serovars found in our study may suggest exposure of dogs to pathogenic *Leptospira* from multiple sources. Previous studies on Amazonian wildlife have shown a high diversity of *Leptospira* serovars [21,22]. Therefore, serovars may not be as species-specific as currently thought and wildlife might be an important source of serovar diversity which, due to the nearness to the forest, likely influences the ecoepidemiology of leptospirosis in the studied Kichwa communities.

It is important to note that none of the dogs in our study were vaccinated against leptospirosis, and the high diversity of serovars encountered suggests that even vaccination, as routinely performed in Ecuador and other Latin American countries, would have little effective in preventing canine infection or carriage, although it may reduce the prevalence of certain serovars. In Ecuador, the health authority, AGROCALIDAD, has registered and approved multiple vaccines for canine leptospirosis [78]. Most of them are bivalent and include serovars Canicola and Icterohaemorrhagiae, but there are also two approved multivalent vaccines that might have higher coverage (one contains serovars Canicola, Icterohaemorrhagiae, Pomona, and Grippotyphosa, and the other contains Canicola, Icterohaemorrhagiae, Grippotyphosa, Pomona, Tarassovi, and Wolffi). Importantly, we found that a high percentage of samples cross-reacted with multiple leptospira serovars (44.4%). This has been commonly reported in samples from dogs in the acute phase of the disease or due to common leptospiral antigens in other serovars [79,80]. Common antigens across serovars can also result in attribution of a sample to the wrong serovar. The problems can be overcome by using local isolates that have been previously characterized and whose cross-reactivity is known [80]. However, few efforts have been made in Ecuador to obtain, culture, maintain, and test local *Leptospira* isolates.

Dogs infected with the pathogen will begin to excrete bacteria in their urine 7 to 10 days post-infection, and this excretion can continue for several weeks or even years [37,81]. Additionally, asymptomatic shedders have been found in 0.2 to 48.8% of dogs worldwide [82,83], but such studies are rare and much is unknown about how frequently this occurs . Surprisingly, our results show that 94.7 % (n=19 - CI: 73.9-99.8) of the asymptomatic dogs were excreting *Leptospira*, the highest percentage reported so far. Three pathogenic *Leptospir*a species were identified in dog urine, *L. santarosai*, *L. noguchii*, and *L. interrogans.* These species have been previously identified in South America [84–86]; *L. santarosai* and *L. noguchii* have been reported in asymptomatic wildlife and dogs, and all three have caused severe disease in humans [22,28,83,87–95]. *L. interrogans* is the most common and widely distributed pathogenic species and known to also infect rodents and small mammals [96–98]. In our study, *L. interrogans* was found only in a community close to the small town of Pompeya, although the presence of this species in more remote communities cannot be excluded due to the small number of samples collected. *L. noguchii* and *L. santarosai* were found in remote communities. The relatively high genotype diversity is consistent with the high serotype diversity and given the close interaction of dogs with the environment and wildlife, is suggestive of varied sources such as might be encountered in the adjacent forest.

The methods used to preserve and transport samples, coupled with detection and sequencing methods, reduced the likelihood of false positives. Recovering *Leptospira* DNA from urine is complicated due to pH and degradation of leptospiral DNA at ambient temperatures or after freezing and thawing as might occur during transit [99]. In the Amazon basin, consistent cold storage of samples is not possible. We therefore used RNA/DNA shield (Zymo) to preserve DNA integrity. We were able to increase the sensitivity of *Leptospira* DNA detection by combining the results of two highly sensitive TaqMan assays capable of detecting 1×10^1^ copies/µL [17,51]. By using both assays, we reduced the likelihood of false negatives and doubled the number of positive samples.

Researching leptospirosis within the framework of the One Health concept [100] by considering the particularities of the disease in different settings is essential. In rural areas, we expect more complicated transmission and cycling networks because of high genetic heterogeneity of *Leptospira* and interactions with diverse host species [2,17,29,101]. In the Amazon region of Brazil, Peru, and Bolivia, wild animals like *Marmosa* spp., coati, nine-banded armadillo, opossum, porcupine, rodents, primates, bats, and wolves are exposed to the disease or excrete leptospires in their urine [21–25,102]. Details about the pathogen and host diversity, density of animal reservoirs, abiotic factors, and interactions among all these elements will ultimately guide the design and implementation of effective prevention and control plans. Our results add important information to existing general knowledge by suggesting that dogs might not only be an important risk factor for human leptospirosis, but also might contribute to the sylvatic transmission cycle. Certainly, much remains to be learned about the epidemiology of leptospirosis in a megadiverse place like the Amazon basin. Undoubtedly, a major challenge to understanding disease cycling in one of the world’s most diverse ecosystems will entail the logistical and methodological hurdles of sampling wild animals.

## Conclusions

A high level of seropositivity and prevalence of pathogenic *Leptospira* DNA in dogs from five indigenous Kichwa communities provides strong evidence that infection and carriage of leptospirosis is very common among dogs in the Ecuadorian Amazon basin. The high serotype and genetic diversity of samples, coupled with the lack of a single dominant type suggest that there may not be any serotype of genotype that is specifically adapted to dogs, and sources of transmission to these dogs are likely to be varied. The domestic dogs in these communities are free-roaming and often hunt wildlife and interact with the ecosystem of the adjacent forest, providing frequent opportunities for the transmission of *Leptospira* to and from wild animals. Importantly, frequent interactions with humans and presence in the peridomestic space is likely to result in transmission between humans and dogs, and the high prevalence in dogs suggests that human leptospirosis in this region is likely greatly underestimated. To more completely understand the cycling and transmission of *Leptospira* in this environment, high-resolution genotyping of longitudinal samples collected from dogs, humans, wildlife, soil, and water would be ideal. This work however establishes that domestic dogs are likely to play an important role in leptospirosis epidemiology in this region, with implications in other regions of the world where peridomestic animals interact with surrounding environments and wildlife.

## Acknowledgments

We would like to acknowledge the five Kichwa communities in northern Yasuní National Park, Centro de Bioinformática – Universidad San Francisco de Quito, Cristopher Pineda, and Andrea Guayasamin.

---

**S1 Table.**
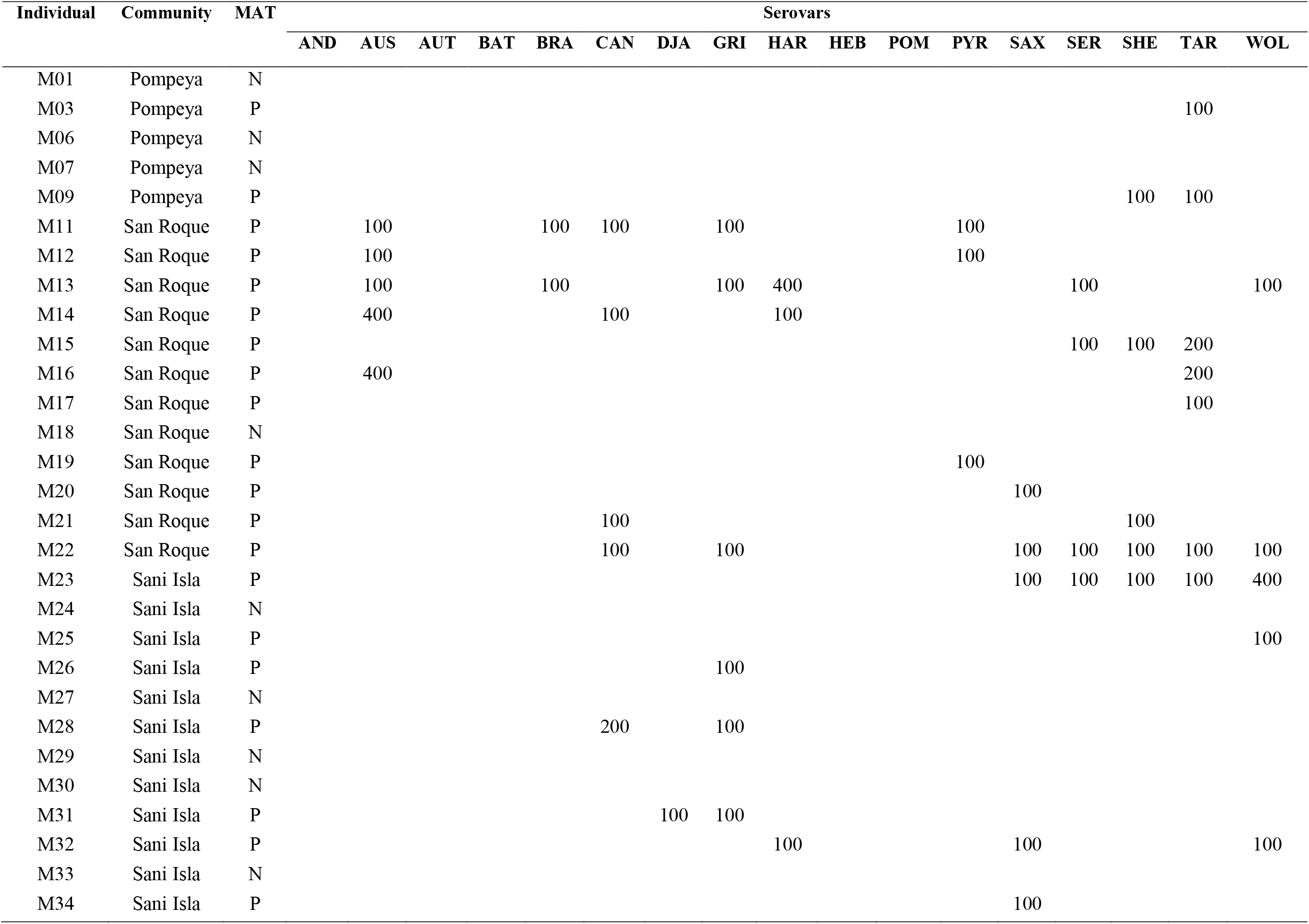
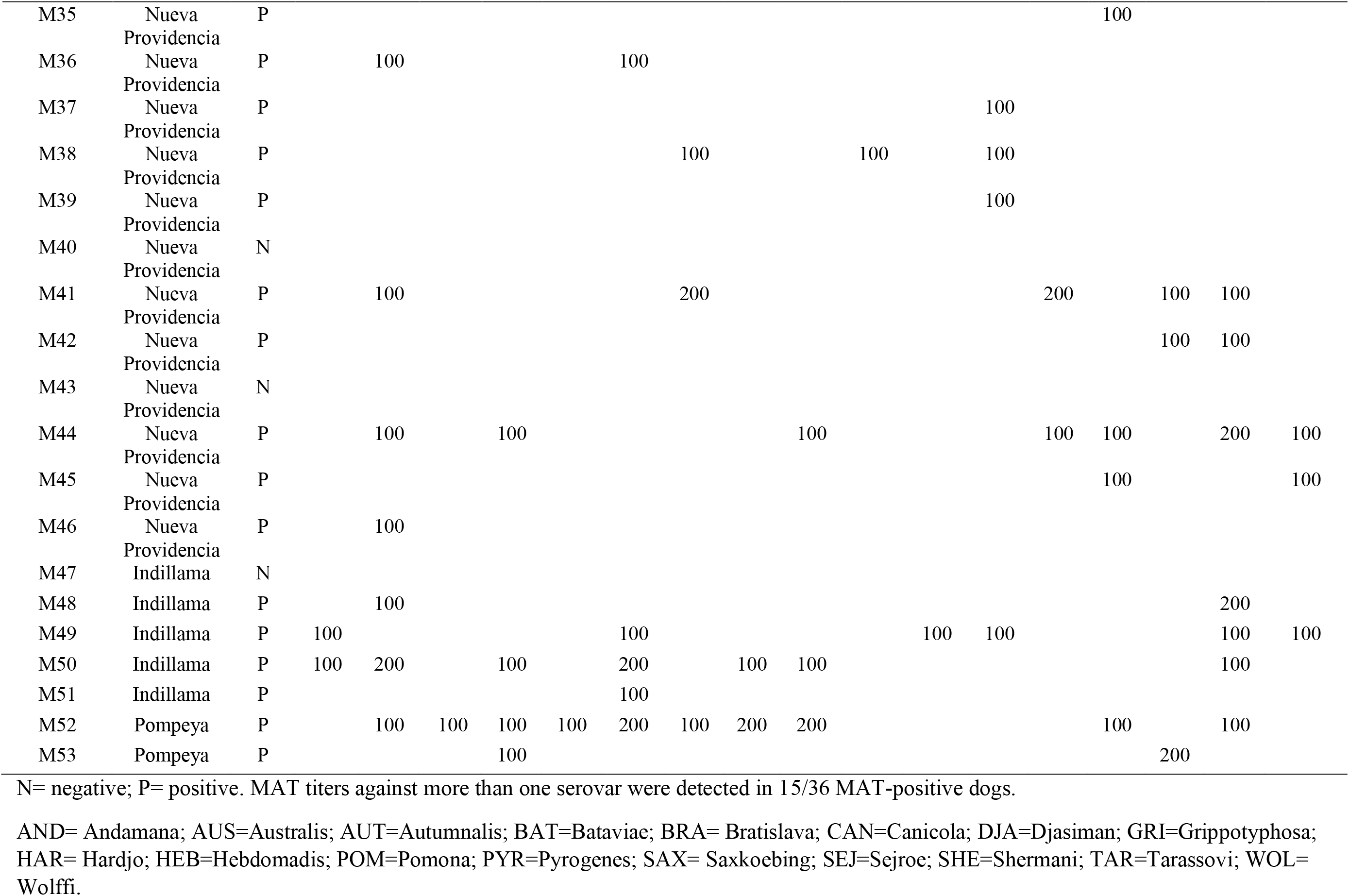
Serological analysis of 48 dogs by MAT.

